# A real-time multiplexed microbial growth intervalometer for capturing high resolution growth curves

**DOI:** 10.1101/533356

**Authors:** David C. Vuono, Bruce Lipp, Carl Staub, Evan Loney, Joseph J. Grzymski

**Author notes:** **Correspondence:** Joseph J. Grzymski.

## Abstract

Batch cultures are a low maintenance and routine culturing method used in anaerobic microbiology. Automated tools that measure growth curves from anaerobic microorganisms grown in traditional laboratory glassware, such as Balch-type tubes, are not commercially available. Here we present a new MicrobiAl Growth Intervalometer (MAGI) that captures microbial growth curves through photo conductivity of the medium using a diffused light pattern of specified frequency, rather than photo-attenuation of collimated light used in traditional systems, and is configured with an offset photodetector/emitter to minimize direct impingement of light from the source to improve the resolution of the solution’s density. MAGI is operated by software-driven automation and offers investigators a low noise/high gain instrument with capabilities for remote visualization and data acquisition. MAGI is a low maintenance, low cost, and robust platform primarily for anaerobic cultivation and growth monitoring. We demonstrate the utility of this device by first showing that growth rates and generation times in *Escherichia coli* K-12 are reproducible to previously published results. We then tested MAGI to measure growth curves of an environmental organism, *Intrasporangium calvum*, under various media compositions. Our results demonstrate that MAGI is a versatile platform to measure growth curves in media under various redox conditions (microaerobic and anaerobic), complex mediums such as Luria-Bertani (LB) broth and minimal media, and for resolving diauxic growth curves when *I. calvum* is grown on a disaccharide. Lastly, we demonstrate that the device can resolve growth curves for μM concentrations of resources that yield low biomass. This research advances the tools available to microbiologist aiming to monitor growth curves in a variety of disciplines, such as environmental microbiology, clinical microbiology, and food sciences.

## 1 Introduction

Microbial growth and physiology investigations rely on two cultivation techniques: batch cultivation (Monod, 1949) and continuous cultivation, such as the chemostat (Monod, 1950). Both culture systems are designed to produce cells as a function of specific growth rate (μ, h^−1^). In continuous cultures, μ is set *a priori* by the investigator through a predefined dilution rate of fresh incoming media into the culturing vessel, where the cells are held in a steady-state of nutrient limitation. This state of nutrient limitation is equivalent to the cusp of the growth curve at late-exponential/early-stationary phase for microorganisms growing in the same media in batch cultures that have nearly exhausted the same growth limiting nutrient (Saldanha et al., 2004). Both batch and continuous culture techniques are complementary tools in a microbiologist’s toolbox and are used to investigate the influence of μ on microbial physiology and *vice versa* (Egli, 2015). The chemostat allows investigators to focus and study microbial physiology continuously at a single growth phase (late-exponential/early-stationary) and predefined growth rate. Whereas batch systems allow investigators to prepare media of various nutrient compositions to observe all growth phases (lag, exponential, stationary, and death) and investigate the effects of nutrient composition on growth rate in parallel (Ingraham et al., 1983).

Both culturing techniques are routinely used for aerobic and anaerobic microbiology but batch systems are better suited for screening cells for growth under a multitude of nutrient compositions and environmental conditions simultaneously. Many high throughput technologies are available to screen cells for growth under aerobic conditions, such as plate readers, however these screening tests are performed in small volumes (μl scale) making most omic-based investigations on the same biological material unfeasible. Furthermore, there are no commercially available high throughput and automated technologies for screening anaerobic cultures in parallel in standard laboratory test tubes and many do-it-yourself (DIY) based systems (Maia et al., 2016; Sasidharan et al., 2018; Takahashi et al., 2015) require personnel time that investigators must allocate towards construction and troubleshooting rather than research. Many of these systems are also not easily scalable to increase experimental replication. As far as standard laboratory glassware, Crimp-sealed culture tubes, such as Balch tubes, are routinely used in anaerobic microbiology for culturing microorganisms in larger volumes (e.g., 10 - 20 ml). However, generating growth curves from these culture vessels typically requires manual measurements, using optical density (OD) or gas production measurements taken at regular intervals (Bräuer et al., 2006), and these measurements can be laborious and time consuming (in high replication) especially for slow growing organisms.

To fill this technological void, we developed a high-resolution MicrobiAl Growth Intervalometer (MAGI) (i.e., capturing optical density data at predefined intervals) capable of multiplexing biological replicates in standard laboratory glassware. MAGI is unique from traditional OD spectrophotometers in several ways. Traditional OD spectrophotometers operate on the principle of photo attenuation where collimated light, either a broadband light source passing through a pass filter of a specified wavelength or a light source with specified frequency, passes through a medium to a photodetector directly opposite the light source (Matlock et al., 2011). The amount of light reaching the photo detector is then proportionally attenuated (i.e., scattered) by biomass in the media and the signal at the detector is reduced. An inverse function is then applied to the data so the investigator can observe the growth curve in the traditional shape (lag, log, stationary, and death). In MAGI, growth monitoring is based on photo conductivity where a diffused light source of specified frequency (or frequencies) is used to illuminate the test tube. The photo-detector is then offset from the emitter at a distance sufficient to minimize direct impingement of light from the source onto the photo-detector and the amount of light arriving at the photo-detector increases proportionally by photo conductivity of the medium resulting in an increased signal.

MAGI is compatible with traditional Balch tubes common to all microbiology labs. Standard preparation techniques, such as Hungate technique, are used to dispense media into the culture tubes and once sealed with butyl-rubber stoppers and crimped, the investigator can inoculate the various medias with the target organism, insert the vials into the MAGI device, and autonomously measure accurate low-noise growth curves in real-time at the click of a button. MAGI uses software-driven automation to operate and collect OD data that is hosted on a server and can be visualized and acquired remotely. MAGI is constructed around a robust optical device and circuit board (light emitter/detector and amplifier gain station, Figure 1) and inexpensive, robust, and commercially available automation hardware (Arduino microcontrollers, raspberry pi computer). This configuration enables a low construction cost that can be passed onto consumers to affordably obtain high resolution growth curves of anaerobic microbial cultures. The device implements 12 parallel culture tube slots, with capacity for upgrades that are constrained only by the size of the incubator/shaker available to the investigator. We use open-source computer languages written in Python, bash, and Arduino (C++) to operate MAGI. Data capture intervals are set through a user-defined programable function.

**Figure 1.**
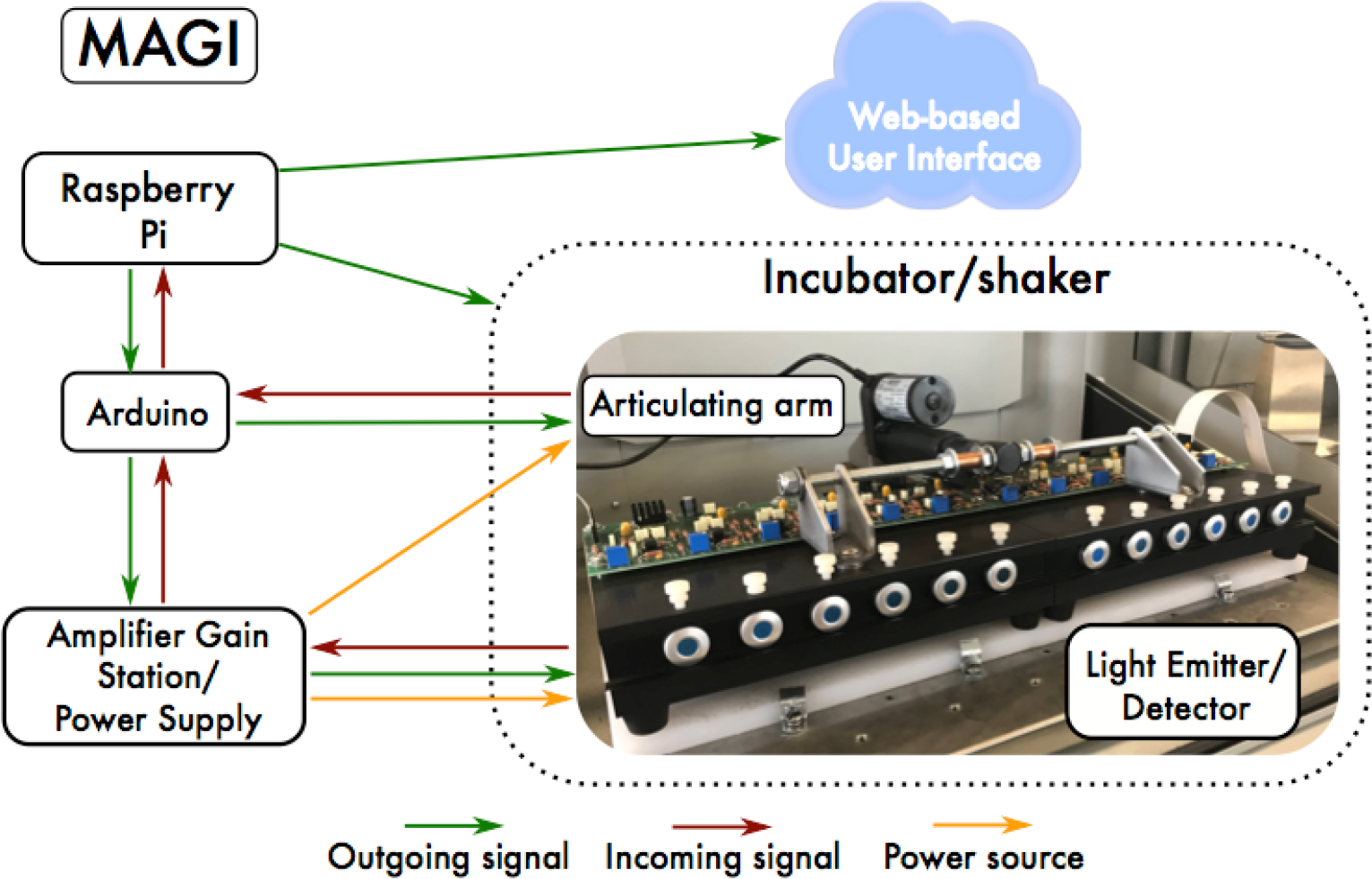
Schematic of MAGI. The Raspberry Pi is a $35 USD computer that controls all of the hardware devices needed to collect growth data from anaerobic organisms. Collected data is configured to be visualized and acquired remotely through a Web-based user interface.

## 2 Materials and Methods

### Media preparation

Preparation of lactate/nitrate media was conducted in a 2L Widdel Flask. After autoclaving, the media was immediately put under an anoxic headspace (N2/CO2 80:20 mix) and sterile filtered (0.2μm) trace elements, trace vitamins, and reducing agent were added. The media was cooled under an anoxic headspace and buffered with bicarbonate to maintain a pH of 7.2. Hungate technique was used to dispense media into standard Balch tubes (18×150-mm glass tube) (20 mL) pre-flushed with a sterile stream of ultra-high purity (UHP) N_2_ and sealed with blue 1” butyl rubber stoppers. All cultures were grown at 30 ºC and shaken at 250 rpm. Nitrate reducing minimal media was prepared with the following final concentrations: NaCl (0.06 mM), NH4Cl (1.4 mM), MgCl2 (0.2 mM), CaCl2 (0.04 mM), KCl (0.1 mM), K2HPO4 (1.1 mM), NaHCO3-(30 mM), cysteine (1mM) as reducing agent, resazurin as redox indicator, and trace elements and trace vitamin solutions as reported (Hillesland et al., 2014; Yoon et al., 2013). 1M sterile filtered (0.2μm) concentrated stocks of 60% w/w sodium DL-lactate solution (Sigma-Aldrich, St. Louis, MO, USA) were diluted into media prior to autoclaving. Culture tubes were inoculated with cells harvested from a previously grown batch culture in late exponential phase in order to ensure that each experiment began with cells in the same metabolic state.

### Strain information

For growth experiments, we used the Invitrogen OneShotTOP10 Electrocomp *Escherichia coli* K12 strain and *Intrasporangium calvum* strain C5, which was isolated from a nitrate contaminated groundwater well at the Oak Ridge National Laboratory, Oak Ridge, TN, (Lat: 35.97990, Long: 84.27059) as reported by (Vuono et al., 2019).

### Data processing and operational software

Data output from MAGI is an analog-to-digital (ADC) conversion. Because spectrophotometric measurements are relative, and must be constrained to a quantitative measure through standard curves, such as cell count. However, for the purposes of our work (to demonstrate the functionality of MAGI), we chose to report our biotic results as OD_600_, which is traditionally used by microbiologists. Conversion from ADC units to OD_600_ was achieved by collecting a starting and ending OD_600_ value for each culture tube (Hach DR2500 Spectrophotometer). Blanks, specific for each media, were used to zero the instrument prior to collecting the starting value. Using the recorded values by MAGI, OD_600_ values were then populated using linear interpolation. However, because OD methods are a spectrophotometric, standard curves of cell concentration are needed to accurately quantify cell concentration. Cell numbers could easily be calculated from the ADC units, as reported elsewhere (Vuono et al., 2019). However, we chose to present our data as OD_600_, since it is commonly used by microbiologist to measure growth curves.

Growth rates were calculated using the package Growthcurver (Sprouffske and Wagner, 2016) in the R environment for statistical computing. The software used to operate MAGI is open source and available on Github (https://github.com/dvuono/MAGI)

## 3 Results and Discussion

### Operation

MAGI is a collection of hardware devices consisting of an Arduino Uno microcontroller with an ethernet shield, a raspberry pi computer running Linux Ubuntu, an articulating arm (powered by a linear actuator for raising the culture tubes in a vertical position), a power source with analog-based signal amplification for each measurement channel, and a multiplexed light emitter/detector that houses the test tubes (Figure 1). In simplest terms, MAGI runs software to interact with hardware at predefined intervals (i.e., intervalometer). An intervalometer is a device that counts intervals of time. Such devices are commonly used to signal, in accurate time intervals, the operation of some other device. For instance, an intervalometer might activate a device every 30 seconds. These intervals are triggered by Python scripts based on a raspberry pi computer, which communicates primarily with an orbital shaker/incubator and the articulating arm. The timing of the sampling interval is defined by the user through the software utility cron, which is a time-based job scheduler in Unix-like computer operating systems. People who set up and maintain software environments use cron to schedule jobs (commands or shell scripts) to run periodically at fixed times, dates, or intervals. Cron is most suitable for scheduling repetitive tasks. Cron is driven by a crontab (cron table) file, a configuration file that specifies shell commands to run periodically on a given schedule. Once the sampling interval is defined, MAGI first communicates with the orbital shaker/incubator through ASCII commands sent via RS-232 serial commands. The command syntax varies between manufacturers; however, this information is readily available in user manuals provided by manufacturers.

The operation of MAGI consists of five basic operational steps: First, communication through software on the raspberry pi signals the stop command of an orbital shaker/incubator (used to incubate and mix microbial cultures) through a standard serial communication channel. Second, once shaking has stopped, 5.0 volts (V) of direct current are sent from the Arduino to activate the articulating arm that raises the light emitter/detector in the upward position to orient the tubes vertically. Third, optical density data is collected after a short delay period (to allow bubbles to rise to the surface of the medium) and recorded at user specified time intervals. Fourth, the 5.0 V circuit to the articulating arm is interrupted, which lowers the light emitter/detector. Fifth, the raspberry pi signals the start command through serial communication channel to resume shaking of the orbital shaker until the next sampling time. This process repeats until the investigator terminates the experiment.

Each data series is time-stamped and populated in tab-delimited format on the Raspberry Pi. Data from each successive sampling interval is appended to the text file until the experiment is terminated. Meanwhile, the contents of the data file are displayed in a graphical format on a web page hosted on the Raspberry Pi. The web page is coded in HTML and Javascript to provide the investigator with an interactive experience. A simple interface allows the investigator to monitor the progress of the experiment or export and save the data for external use. At the end of the experiment, the data file may be cleared in preparation for additional experiments.

### Technical Specifications

Replication is essential in microbiology because growth and other physiological responses are sensitive to variations in culture media, preparation techniques, and growth conditions (Egli, 2015). Chemostats are typically expensive ranging from $20-30K where replication in parallel is often not feasible due to cost. Maintaining fresh media stock for continuous operation is also laborious. Thus, replication in continuous culture is usually performed sequentially through time, or serially, which leads to dependencies of autocorrelation and hysteresis. Batch experiments, however, can be multiplexed to accommodate as much reproducibility as is needed by the investigator. For MAGI, we implemented 12 culture tube measurement cells to support adequate reproducibility for experimental and control groups. The insert size for each measurement cell is designed to fit standard laboratory glassware, such as Balch-type tubes, with dimensions 18mm x 150mm.

The light detectors are silicon-based, shielded, and hermetically sealed quartz windows, which enable wide band sensitivity. For the light emitter, light emitting diodes (LEDs) of 618 nm were chosen to avoid absorption maxima of resazurin at 600 nm (Venkata Rami Reddy and Bordekar, 1999), a common redox indicator used in anaerobic microbiology. The light emitter consists of LEDs that have a nominal dominate wavelength of 618 nm with ±11.5 nm full width at half maximum (FWHM), a peak wave length of 624 nm, ±30º angle of half intensity, and 90% of total flux is captured within 75º. This creates an emitter with a diffused light source, rather than collimated light (often used in traditional OD spectrophotometers), and is driven by 65 mW of constant energy. Traditional spectrophotometers typically operate through photo-attenuation of collimated light, where the detector is located directly across from the emitter (Koch, 1970). In MAGI, the emitter is positioned at the base of the culture tube and is intentionally offset from the detector (Figure 2). The emitter and detector are offset because this increases the pathlength the light must travel, thereby allowing the microorganisms to function as a photo-conductive medium and increase the resident resolution of the solution’s density. This configuration requires significantly more amplifier gain (*A_V_*) and is achieved using a transimpedance amplifier to permit low noise/high gain for each measurement. The photodetector system is thus comprised of a photodiode operating in transimpedance mode with an amplifier, which is then tunable by the investigator through adjustments of the base voltage at the amplifier gain station (Figure 1). As is standard in most optical density devices, voltage at the photodetector decreases as cell number (optical density) increases and this analog signal is converted to a digital unit (analog-to-digital conversion; ADC). The ADC unit can then be readily converted OD_600_ by recording the starting and ending optical density of the culture tube and interpolating the OD_600_ values based on recorded ADC values.

**Figure 2.**
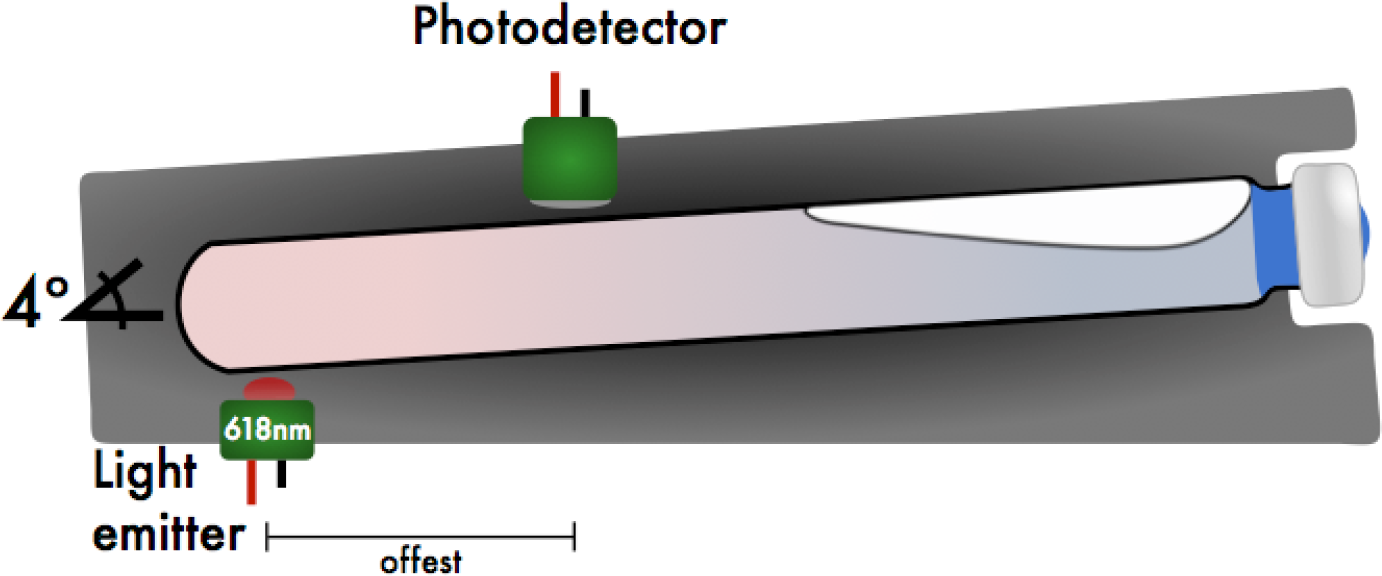
Sagittal view of the light emitter/photodetector and culture tube position. High resolution of MAGI is achieved by off-setting the light emitter from the photodetector. A default tube angle of 4º is implemented to help migrate bubbles produced from biosurfactants towards the top of the culture tube as the instrument stops shaking, while still maintaining a horizontal position while shaking.

### Countering Optical Interference from Foaming and Biosurfactant Production

To validate the need for control from biosurfactant interference on optical readings, we simulated bubble formation in culture tubes (n=12) using standard laboratory soap (0.1% v/v Liquinox). Interference of the optical path from surfactants was tested at 1-minute sampling intervals under two states where readings were taken with 1) culture tubes kept at the default horizontal position between and during optical readings (4º Tilt) and 2) with culture tubes lifted by an articulating arm to a 30º angle relative to horizontal during optical readings (30º Tilt) (Figure 3). Initially, the first two readings were taken with the tubes oriented at a 30º tilt to define a baseline of 182.1 ± 11.2 ADC units. In between the 1- and 2-minute readings, the lead was removed from the 5.0 V relay on the Arduino to prevent activation of the articulating arm. This allowed for optical readings to still be taken but at the horizontal position (4º tilt) where the optical path to the light detectors would be occluded by bubbles at the liquid-air interface. Upon the first reading, the optical signal increased immediately to 955.2 ± 90 ADC units and remained elevated for the duration of position, with an average of 994.1 ± 53.4 ADC units. After the last reading with a 4º tilt (22 minutes), the lead was reconnected to the 5.0 V relay on the Arduino to engage the articulating arm. This resulted in the optical signal dropping back down to baseline for the remainder of the experiment, with an average of 182.2 ± 11.5 ADC units. These results demonstrate the importance of controlling for biosurfactant interference to prevent aberrant signals.

**Figure 3.**
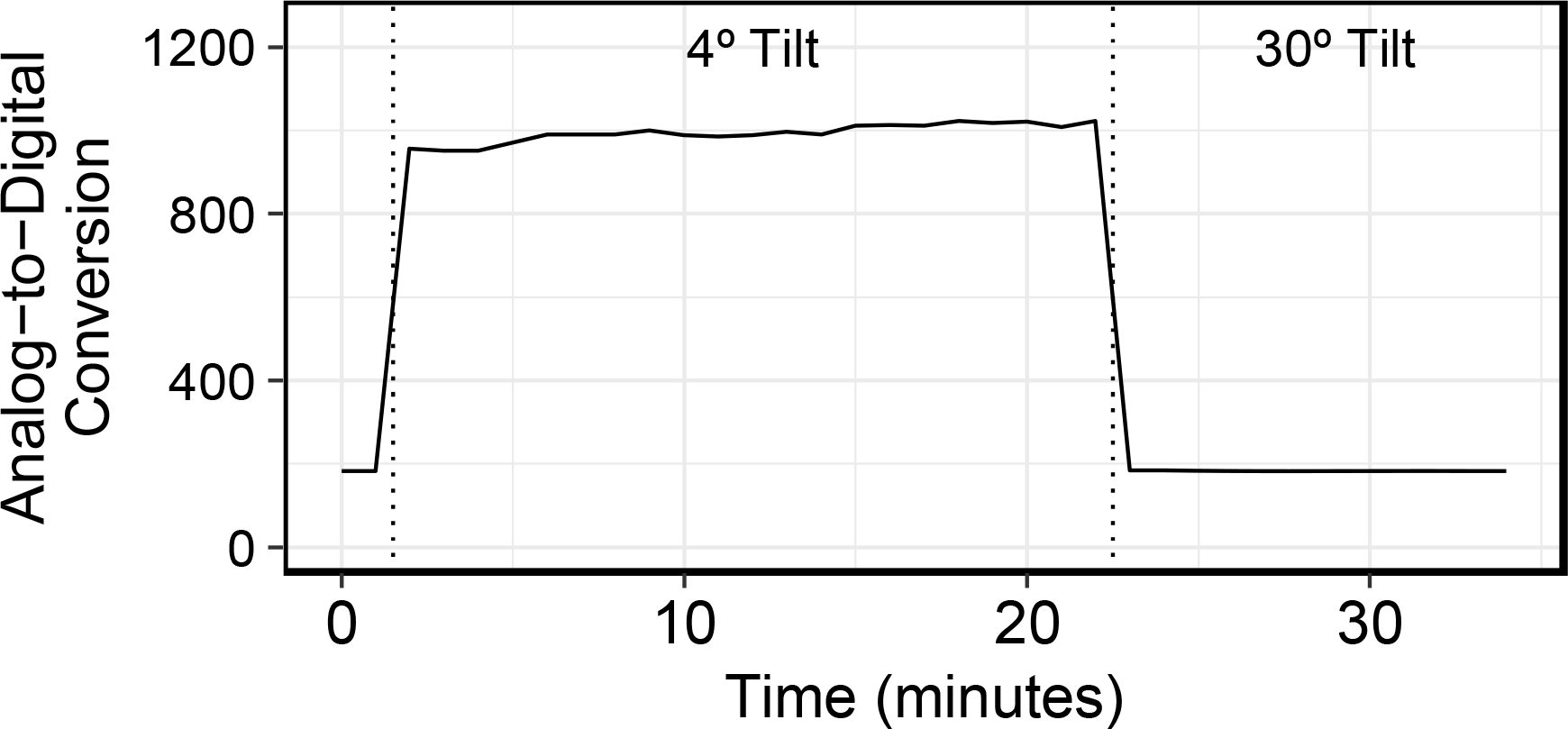
**(A)** Effectiveness of an articulating arm to orient tube position to a 30º tilt during data acquisition to obviate biosurfactant obstruction of the photodetector.

### Capturing Growth Curves from Laboratory and Environmental Organisms

To test the effectiveness of MAGI at measuring batch growth of laboratory organisms, we selected *E. coli* K-12 for growth in autoclaved LB medium that was prepared aerobically and dispensed in Balch tubes capped with butyl rubber stoppers. This design enabled us to capture growth curves from aerobic *E. coli* cultures in a sealed culture tube compatible with horizontal shaking. The OD sampling interval was 6 minutes in order to capture the exponential growth phase of *E. coli*. Results show that MAGI was able to capture a typical microbial growth curve from lag phase to stationary phase of *E. coli* grown aerobically in LB media (Figure 4A). The transition to stationary phase was observed as an abrupt transition due to the optical density within the culture vessel reaching the maximum resolution of the light detector. However, this abrupt transition did not prevent standard microbial growth parameters to be calculated: the specific growth rate (μ) was 2.0 ± 0.05 h^−1^ with a generation time of 20.7 ± 0.5 minutes (Figure 4A Inset). These results are congruent with values reported elsewhere (Berney et al., 2006), thereby demonstrating the ability of MAGI to accurately measure growth curves by which standard microbial growth parameters can be calculated.

**Figure 4.**
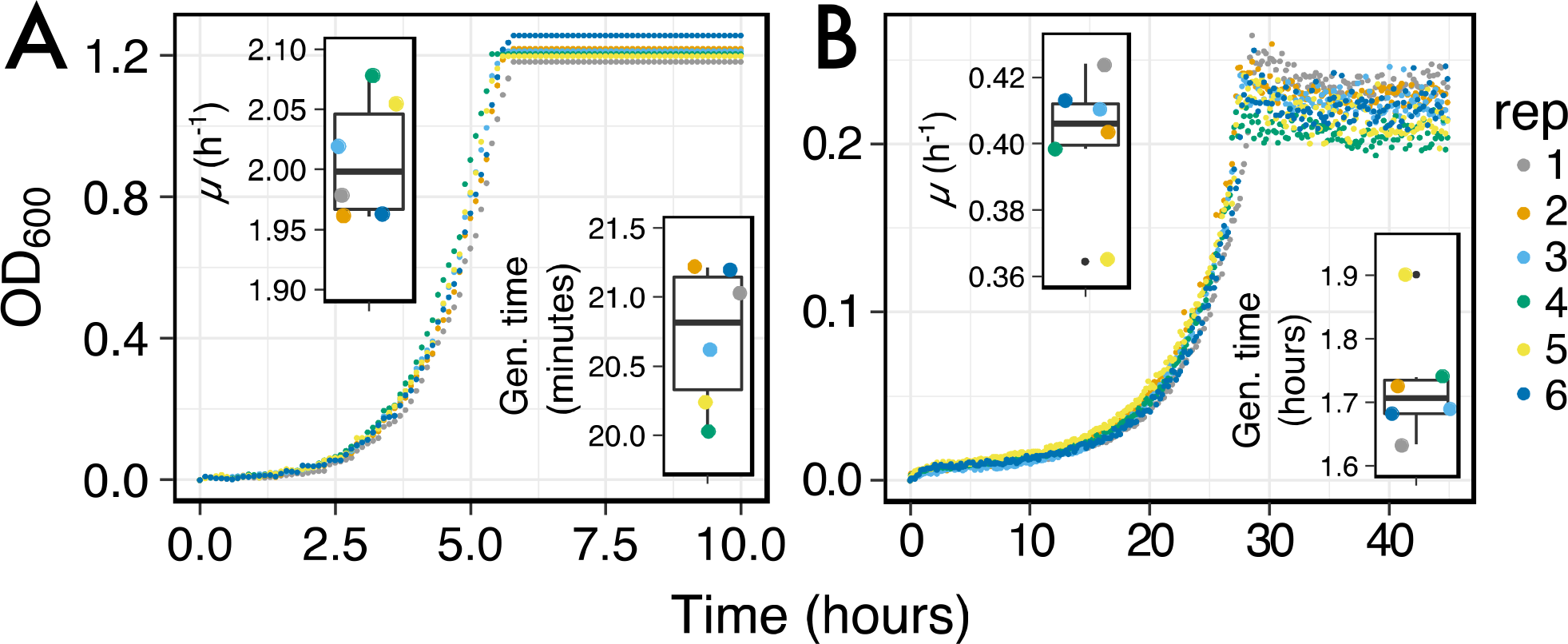
Growth curves of (A) *E. coli* grown in aerobically prepared LB media and (B) *I. calvum* in aerobically prepared minimal media (8 mM Lactate). Inset graphs are of growth rates (top left) and generation times (bottom right) for each respective sample replicate.

Given the reproducibility of growth rate and generation time calculations obtained by the MAGI *E. coli* growth curves, we next generated growth curves from an environmental organism, *I. calvum*, grown with 8 mM lactate as carbon source and electron donor and 3 ml of sterile air injected into the anaerobically prepared vial. *I. calvum* is an Actinobacterium of the family *Intrasporangiaceae* (del Rio et al., 2010). It known for producing branching mycelium with intercalary sporangia in the mycelial hyphae (Kalakoutskii et al., 1967), the function of which is unresolved (Lechevalier and Lechevalier, 1969). *I. calvum* has been isolated from air (Kalakoutskii et al., 1967), terrestrial subsurface (Green et al., 2010; Vuono et al., 2019), has been found in activated sludge bioreactors (Vuono et al., 2015), and is known for its ability to dissimilate nitrite via a dual-pathway of denitrification and ammonification (Vuono et al., 2019).

When *I. calvum* was grown with lactate and O_2_, we observed growth curves where all replicates were in sync during each growth phase (Figure 4b). Similar to the *E. coli* growth curves (Figure 4A), *I. calvum* displayed a similar abrupt transition to stationary phase. However, this was likely due to the depletion of O_2_ in the culture vessel rather than maxing out the voltage differential between emitter and photodetector. From these growth curves, we were able to calculate a specific growth rate (μ) of 0.4 ± 0.02 h^−1^ and a generation time of 1.73 ± 0.09 hours (Figure 4B Inset).

Next, we asked whether we could measure *I. calvum*’s growth curves from a variety of media compositions in order to resolve minor differences in growth depending on variations in starting conditions. The media compositions tested were microaerobic versus anaerobic (Figure 5A), complex media (Figure 5B), and growth with a disaccharide in order to capture diauxic growth (Figure 5C and D). We prepared anaerobic minimal media using lactate as electron donor and nitrate as electron acceptor (Figure 5A). We then aerobically transferred cells to one culture tube to promote microaerophilic conditions (blue curve/circles, Figure 5A). Microaerobic conditions were verified by visualizing the media turning slightly pink due to the oxidation of the redox indicator resazurin. The second culture tube was inoculated anaerobically where the injection needle was gassed with sterile N_2_ prior to inoculation (orange curve/triangles, Figure 5A). These results showed that MAGI was able to resolve a slightly higher growth rate in the microaerobic culture (Figure 5A, inset top right; μ = 0.2 h^−1^) as compared to the sample that was maintained under anaerobic conditions (Figure 5A, inset top right; μ = 0.18 h^−1^). The cultures growing in complex media (LB) under microaerobic conditions (inoculated similarly to blue curve/circles in Figure 5A) showed a low deviation in growth rates (μ = 0.24 ± 0.004 h^−1^) indicating the accuracy of MAGI to resolve growth curves in complex media (Figure 5B). When *I. calvum* was grown on the disaccharide sucrose, we tested whether the sampling resolution of MAGI could capture growth variation during the rapidly changing biomass phase of exponential growth. For the cultures grown under microaerobic and anaerobic handling (Figure 5C) we observed the characteristic shoulder (short lag period) of diauxic growth (Monod, 1949) at approximately 20 hours (arrows) and were able to resolve the different growth rates during the second exponential phase for each culture tube (Figure 5C, Inset top right; μ = 0.19 h^−1^, 0.12 h^−1^ for microaerobic and anaerobic conditions, respectively). Similar results were observed under aerobic growth conditions (Figure 5D). Overall these results show that MAGI is capable of resolving growth curves from anaerobic and microaerobic growth conditions under a variety of growth mediums.

**Figure 5.**
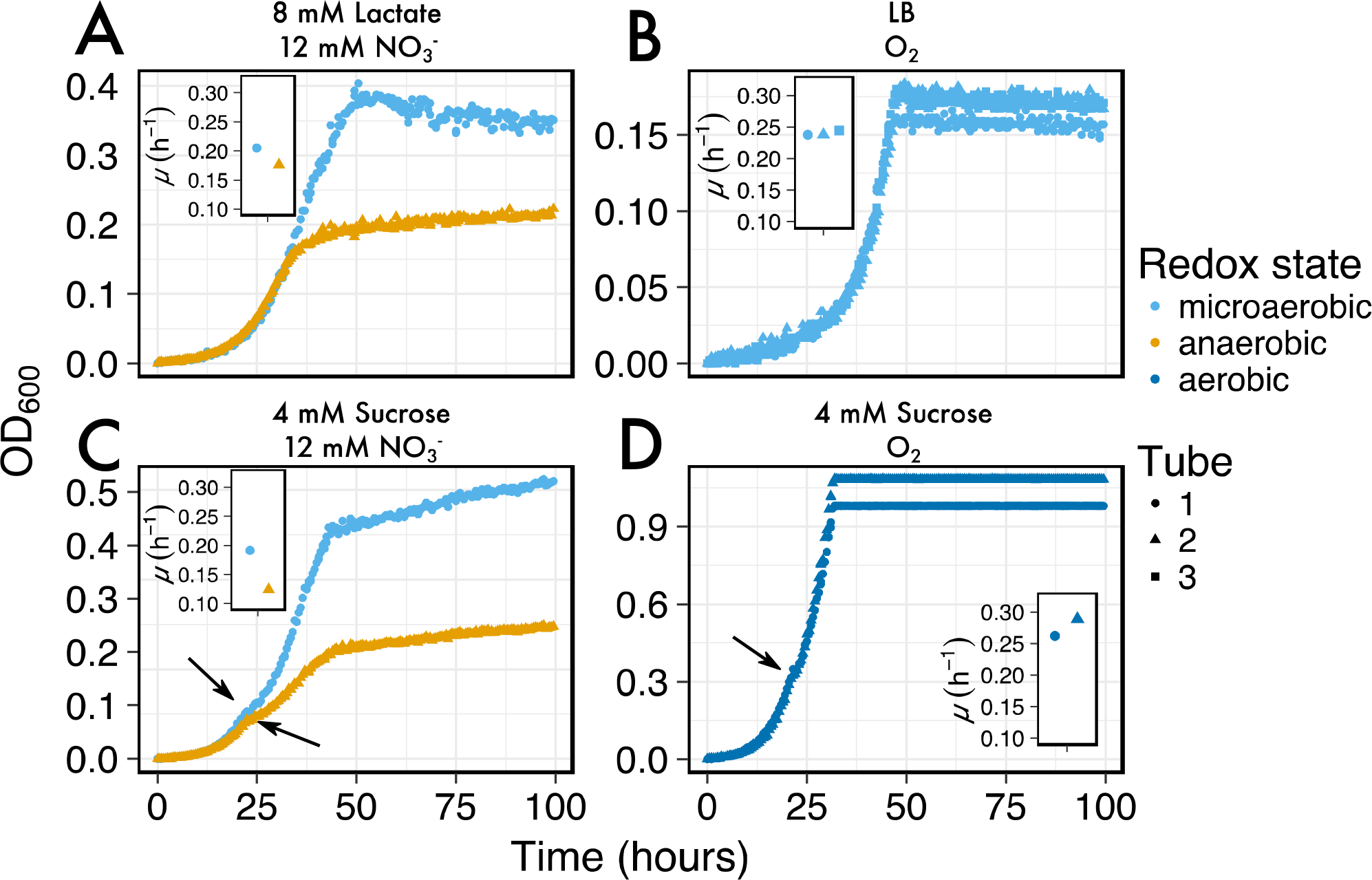
Growth curves of *I. calvum* at various media compositions: (A) microaerobic versus anaerobic, (B) complex media (LB), and growth with a disaccharide under (C) microaerobic versus anaerobic and (D) aerobically prepared media. Arrows in C and D indicate shoulders of the growth curve where diauxic growth transition occurs. Inset graphs are of growth rates for each respective sample tube.

### Capturing Growth Curves from Low Resource Concentration Mediums

Growth yields from anaerobic metabolisms are often times very low due to the lower free energy yields (∆Gº) of electron donors and acceptors used in these reactions (Thauer et al., 1977). Furthermore, measuring growth curves from environmentally relevant resource concentrations (μM amounts) is also difficult due to the detection limits on most OD detection devices. The analog-based signal amplification on MAGI allows users to adjust the detection resolution based on anticipated growth yields. To test this feature on MAGI, we prepared anaerobic media in *I. calvum* cultures with a range of carbon concentrations that varied from mM to μM ranges (Figure 6). Results show that with decreasing carbon concentration, we were able to resolve growth curves to optical densities as low 0.008 OD_600_ (0.04 mM lactate concentration, replicate 3) with group average of 0.015±0.004 OD_600_. These results demonstrate the capability of MAGI to resolve growth curves from media with low substrate concentrations that have low biomass yields. Furthermore, in a previous study we were able to resolve higher growth rates among the treatments (at low carbon concentrations), which selected for the ammonification respiratory pathway over denitrification in *I. calvum* (Vuono et al., 2019).

**Figure 6.**
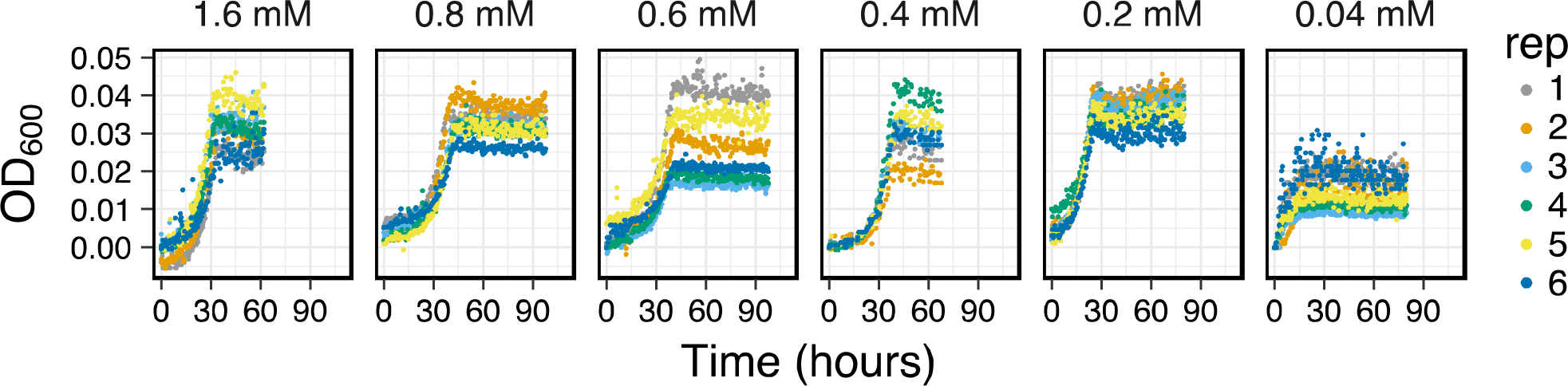
Growth curves of *I. calvum* at various carbon concentrations indicating the sensitivity of MAGI to detect growth curves of cells at low growth yields.

## 4 Conclusion

Batch cultures are a low maintenance and routine culturing method in anaerobic microbiology. Automated tools that measure growth curves from anaerobic microbial cultures grown in traditional laboratory glassware, such as Balch-type tubes, are lacking. Here we present a new microbial anaerobic growth intervalometer (MAGI) that uses software-driven automation, remote visualization, and data acquisition. We demonstrate the utility of this device by first showing that growth rates and generation times in *E. coli* K-12 are reproducible to previously published results. We then tested MAGI to measure growth curves of an environmental organism, *I. calvum*, under various media compositions. Our results demonstrate that MAGI is a versatile platform for anaerobic microbiology research.

## 4 Acknowledgements

We thank N Elliot, DA Stahl, B. Ramsey, A. Murray, Z. Harrold, and P. Longley. A preprint version of this manuscript can be found on the preprint server bioR_X_iv.

## 6 Author Contributions

DV, BL, CS and JG wrote the paper. DV, EL, JG contributed to the conception and design of the study. DV and EL performed lab work. DV, EL, and JG analyzed data and interpreted results. All authors contributed to manuscript revision, read and approved the submitted version.

